# Rapid antibiotic resistance predictions from genome sequence data for *S. aureus* and *M. tuberculosis*

**DOI:** 10.1101/018564

**Authors:** Phelim Bradley, N. Claire Gordon, Timothy M. Walker, Laura Dunn, Simon Heys, Bill Huang, Sarah Earle, Louise J. Pankhurst, Luke Anson, Mariateresa de Cesare, Paolo Piazza, Antonina A. Votintseva, Tanya Golubchik, Daniel J. Wilson, David H. Wyllie, Roland Diel, Stefan Niemann, Silke Feuerriegel, Thomas A. Kohl, Nazir Ismail, Shaheed V. Omar, E. Grace Smith, David Buck, Gil McVean, A. Sarah Walker, Tim E.A. Peto, Derrick W. Crook, Zamin Iqbal

## Abstract

Rapid and accurate detection of antibiotic resistance in pathogens is an urgent need, affecting both patient care and population-scale control. Microbial genome sequencing promises much, but many barriers exist to its routine deployment. Here, we address these challenges, using a de Bruijn graph comparison of clinical isolate and curated knowledge-base to identify species and predict resistance profile, including minor populations. This is implemented in a package, *Mykrobe predictor*, for *S. aureus* and *M. tuberculosis*, running in under three minutes on a laptop from raw data. For *S. aureus*, we train and validate in 495/471 samples respectively, finding error rates comparable to gold-standard phenotypic methods, with sensitivity/specificity of 99.3%/99.5% across 12 drugs. For *M. tuberculosis*, we identify species and predict resistance with specificity of 98.5% (training/validating on 1920/1609 samples). Sensitivity of 82.6% is limited by current understanding of genetic mechanisms. Finally, we demonstrate feasibility of an emerging single-molecule sequencing technique.

## Introduction

The dramatic increase in antibiotic use in healthcare and agriculture since the 1940s has driven a rise in frequency of drug-resistant bacterial strains, which now present a global threat to public health. Clinical isolates resistant to most drugs have now been seen for many species including *Mycobacterium tuberculosis*, *Enterococcus faecium*, *Staphylococcus aureus*, *Klebsiella pneumoniae*, *Neisseria gonorrhoeae*, *Acinetobacter baumannii* and *Pseudomonas aeruginosa*^1^. Antimicrobial susceptibility testing is therefore now central to the treatment of serious bacterial infections diagnosed by culture, and is used to determine the protocols for first-line antibiotic use when culture is not available. At present, phenotyping tests take at least 1-2 days to complete for rapidly growing bacteria such as *S. aureus*, and can take weeks in slow growing bacteria such as *M. tuberculosis.*

Microbial genome sequencing has the potential to substantially increase the speed of antibiotic resistance detection for many pathogens^2^ and in addition provides valuable information on relatedness that could contribute to surveillance. The key biological constraint is the extent of our understanding of the genotype-to-phenotype correspondence – i.e. the genotype needs to be sufficiently predictive of resistance. Increasingly, this correspondence is high for many bacterium/drug combinations. For example, Gordon et al.^3^ recently demonstrated that a curated panel of mutations and genes known to cause drug resistance in *S. aureus* was sufficient to predict resistance for 12 antibiotics with a sensitivity of 97% (95% confidence interval (CI) 95%-98%) and specificity of 99% (99%-100%). Thus at least for one species, a genotype-based method could deliver results with accuracy suitable for use in a clinical laboratory.

Here, we focus on the solving the multiple computational challenges that act as barrier to routine and rapid deployment of such a system in clinical practice. These include not only the need for the tool to be accessible to a user-base who may be unskilled in bioinformatics, but also speed and computational hardware requirements. Finally, and most critically, the system must be validated against the results of existing gold standard testing using extensive and clinically relevant datasets.

In considering these aims, we established two design criteria. First, in order to extend to a wide range of bacterial species, any method would need to be able to detect various categories of resistance-conferring alleles – single-nucleotide polymorphisms (SNPs), indels and entire genes – on potentially diverse genetic backgrounds. The second criterion is a consequence of sample treatment. At present, samples are usually cultured before sequencing; this increases the volume of DNA available for sequencing, at the price of applying a new selective pressure that can result in one strain or species flourishing and dominating the sample. Consequently, any future improvements in sample preparation and sequencing which reduce the culture time towards zero will potentially increase the diversity of the sample. We would therefore want any method to be robust to mixed infections.

Methods using genome sequence data for species identification range from the specific^4^ to the sensitive^5^, but generally performance is measured globally in terms of detection of species presence. However, for clinical use we need considerable flexibility in tuning sensitivity and specificity for different species, potentially weighted to minimize clinical risk. For example, there may be a species associated with high mortality (e.g. *S. aureus*) that can occur in samples mixed with other species (e.g. coagulase-negative staphylococci, which are common contaminants of blood cultures, being present on the skin through which the blood was taken), and that may even share the same resistance genes and thus confound inferences.

Various methods have been used for genotyping resistance features: mutations and genes have been detected by whole genome assembly^3^, genes by assembly and BLAST^6^, or SNPs and indels by mapping^7,8^. These methods have been demonstrated to have adequate performance in some circumstances. However traditional whole-genome bacterial assembly is fundamentally based on the assumption that all data comes from a single haploid genome^9^, and so is ill-suited for mixed samples, and mapping to a single-reference results in error rates that depend on genetic distance of the sample from the reference^10^. There are pre-existing tools for expert users that incorporate resistance prediction^6,11,12^, none of which handle the issue of contaminating related species in clinical samples, or minor clones - we include comparative data below.

Our goal in this study was to move beyond proof-of-concept of how sequencing might work in the clinic, to a general framework for genotype-based antimicrobial resistance prediction, with concrete implementations for two species where drug resistance is of global concern: *S. aureus* and *M. tuberculosis*. We evaluate extensively against clinical gold standards using current (Illumina) sequence data, and demonstrate that the method also works with an emerging strand-sequencing technology (Oxford Nanopore Technologies, MinION). Finally, we will discuss below what is needed to apply our framework to subsequent species.

## Results

### Using population genome graphs for genotyping

We show in Fig. 1a a cartoon of the genetic diversity in a bacterial species and two options for building a reference variation structure. In option i) we show the standard approach, where we select an arbitrary strain (Strain 1) to be the reference genome, along with one copy of each plasmid gene. In this work we have developed an alternate approach, shown in option ii). We start with a curated knowledge-base of resistant/susceptible alleles, and assemble a de Bruijn graph ^13^ of them on different genetic backgrounds, along with many exemplars of resistance genes. This forms our reference graph. In Fig. 1b we show the corresponding analyses of a mixed sample. The traditional approach is shown in option i), whereby sequence reads are mapped to the reference genome and genes, requiring the mapping and inference to cope with the divergence between sample strains and the reference. Our approach (option ii) directly compares the de Bruijn graph of the sample with the reference graph. This results in statistical tests for the presence of resistance alleles that are unbiased by choice of reference or assumptions of clonality. Moreover, these tests will improve as the catalogue of diversity in the species grows. Our approach is implemented in a software framework called the *Mykrobe predictor.* See Methods for details.

**Fig. 1.**
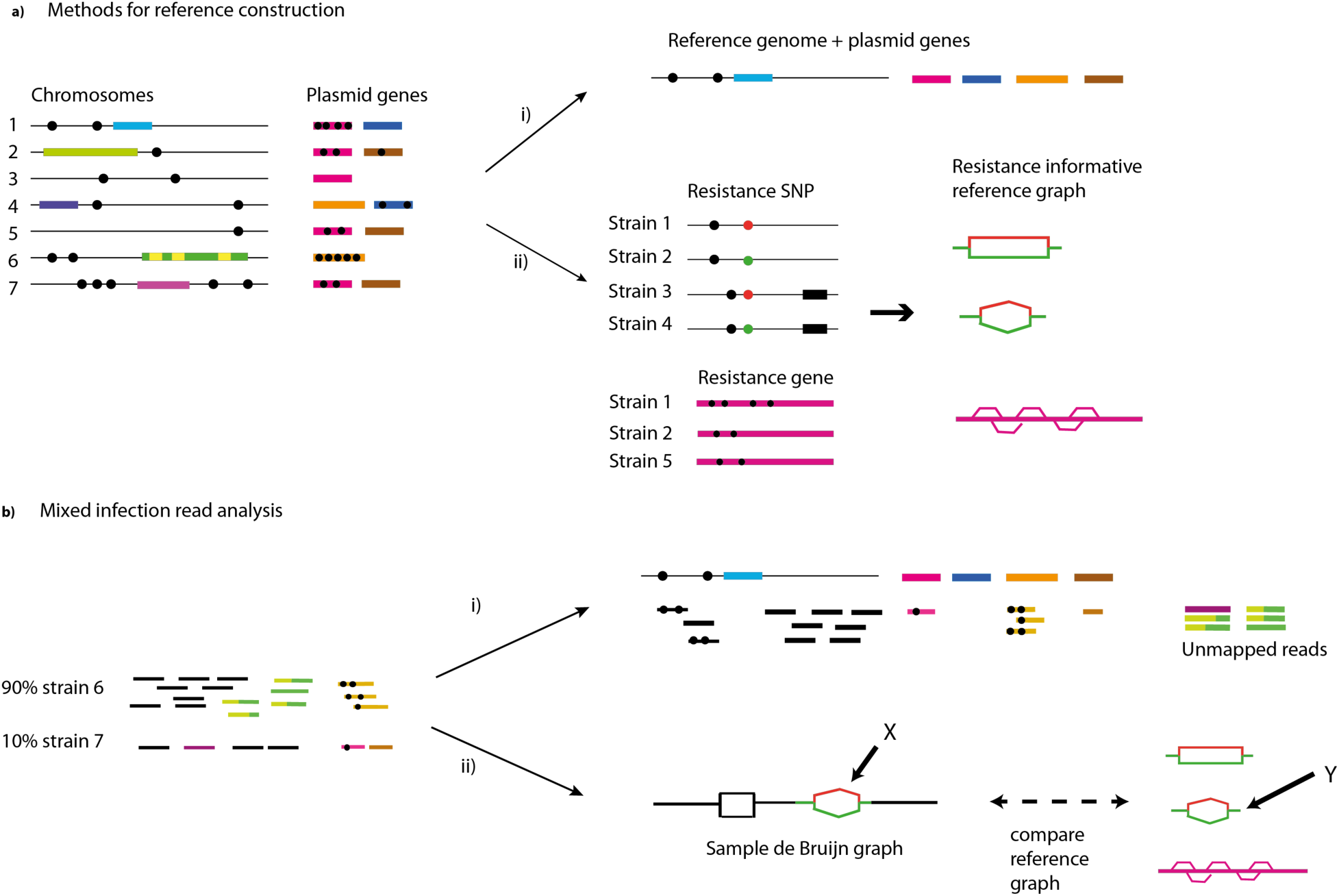
a) Reference construction methods. Left: strains of a bacterial species, showing chromosomes with SNPs (black circles) and genes (coloured blocks). Option i) pick Strain 1 to be reference, plus one example of each plasmid resistance gene. Option ii) our method is to build the de Bruijn graph of all strains, restrict to loci of interest, and annotate resistance (red) and susceptible (green) alleles. For SNPs, local graph topology is determined by adjacent SNPs (black dots) and indels (black blocks). b) Left: sequence data from a clinical sample harboring major (90%) and minor (10%) strains. Right: Option i) map the reads to the reference genome to detect SNPs and genes. Option ii) Our approach: construct the de Bruijn graph of the sample and compare with the reference graph. We see a specific SNP is present both in the sample and the reference graph (marked X,Y). Both the resistant (red) and susceptible (green) alleles are present in the sample, and within-sample frequency is estimated from sequencing depth on each allele.

### S. aureus – species identification

We used dataset St_A (datasets described in Supplementary Fig. 1, Supplementary Table 1), consisting of 532 *S. aureus* and 199 coagulase-negative staphylococci (CoNS) (see Methods, and Supplementary Fig. 1) as a training set to design several panels of probes, used to detect the presence of *S. aureus*, *S. epidermidis*, *S. haemolyticus*, or other coagulase-negative staphylococci. We evaluate our predictions on a separate validation set (St_B) and show the results in Fig. 2a. This confirms an appropriately low rate of missing a true *S. aureus* sample (0/492, upper 97.5% CI 0.7%). We studied the 3 non-*S. aureus* samples that appear to be misclassified as *S. aureus,* and concluded that they were mis-labelled in the NCBI Short Read Archive (SRA), as both BLAST and OneCodex (http://beta.onecodex.com) agreed with Mykrobe predictor. We also created *in silico* mixtures of *S. epidermidis*/*S. aureus* and *S. haemolyticus/S. aureus* (Simulation 1) and obtained 100% power to detect presence of *S. epidermidis and S.haemolyticus* minor infections at frequencies above 0.7% (see Supplementary Fig. 2, Supplementary Methods).

**Fig. 2.**
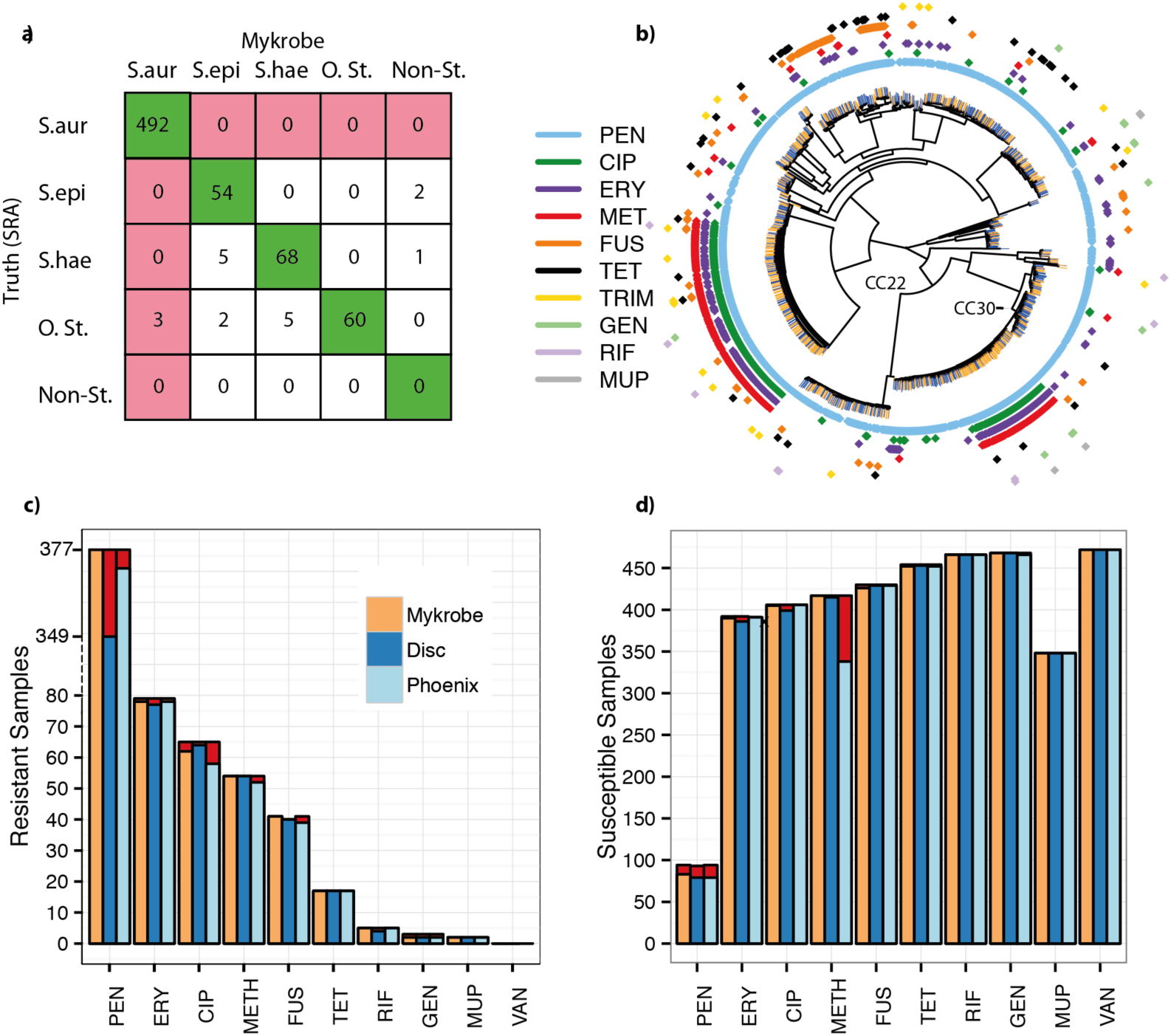
Species and susceptibility predictions for *S. aureus*. a) Species classification results on species validation set St_B. Red shading of box indicates errors we wish to minimize. Abbreviations: S.aur: *S. aureus*, S.epi: *S. epidermidis*, S.hae: *S. haemolyticus*, O.st : other staphylococcus, Non-st: Non staphylococcal. b) Phylogeny of *S. aureus* samples used in evaluating resistance prediction, with tips marked orange/blue to show if sample in training set (St_A1) or validation set (St_B1). Drug resistance in concentric rings around the outside; plasmid-mediated resistance (erythromycin (purple), tetracycline (black)) is distributed across the whole tree. The two multidrug resistant clades are in UK hospital clonal complexes CC22 and CC30. c) Proportion of resistant *S. aureus* samples (St_B1) called as resistant for Mykrobe (yellow), disc test (dark blue) and Phoenix (light blue) compared with consensus, with false negatives in red. Note the break in the y-axis between 80 and over 300 in order to show penicillin on same plot. d) As c) but for calling true susceptible samples as susceptible – false positives in red. A small number of failed disc tests for fusidic acid in panel c) result in a lower bar.

### S. aureus resistance prediction

We use a training set (St_A1) of 495 samples and a validation set (St_B1) of 471 (samples and phenotyping described in Methods, Supplementary Fig. 2). We show in Fig. 2b a phylogeny (construction described in Methods) of these samples, showing that both the training set (orange tips of tree) and validation set (blue) are distributed across the entire phylogeny. We also confirmed also that all major clonal complexes were represented (Supplementary Fig. 3).

All validation samples were phenotyped using two methods: a BSAC disc test^14^ and the Phoenix Automated Microbiology System (BD Biosciences, Sparks, MD, USA), except for trimethoprim, for which only disc testing was performed. A “consensus” phenotype was defined to be that called by disc/Phoenix where they agreed, and the result of an Etest and/or nitrocefin (for penicillin) where disc and Phoenix were discrepant. This allowed us to estimate error rates for disc and Phoenix as well as for *Mykrobe*.

Our prediction algorithm (described in Methods) internally classifies a sample as consisting of a clonal susceptible, a minor resistant or major resistant population, and predicts a resistant phenotype for both the minor and major cases. Note that for drugs where resistance is mediated by genes on variable copy-number plasmids (erythromycin and tetracycline) a minor population with high copy number of a resistance-carrying plasmid may sometimes be called as major resistant. After using the training set data to estimate parameters for our statistical model (see Methods, and Supplementary Table 2) we applied the *Mykrobe predictor* to the validation set. Figure 2, panels c) and d) show in red the false negative calls (panel c) and false positive calls (panel d) for *Mykrobe* and the two laboratory methods: disc and Phoenix. If we consider Fig. 2c first, and focus on the 7 drugs with more than 10 resistant samples, then *Mykrobe* misses fewer resistant calls than the other individual phenotypic methods for all drugs except ciprofloxacin. The FDA requires a diagnostic test have a false negative rate below 1.5% when compared with a reference-lab method^15^. Compared with the consensus phenotype, *Mykrobe* achieved such accuracy for 6 of 7 drugs, with an error rate of 0% for 5 drugs (penicillin, methicillin, fusidic acid, clindamycin and tetracycline), 1.3% for erythromycin and 4.5% for ciprofloxacin. We were unable to determine the reason for the missed ciprofloxacin-resistant predictions, although we note that disc and Phoenix had similar problems, and that this drug has at least one uncharacterized mechanism for resistance^16^. The data underlying this plot are presented in Table 1. The corresponding results for the Disc and Phoenix tests are in Supplementary Tables 3-4.

**Table 1.** *Mykrobe* resistance prediction results for validation set. Resistance prediction results for *Mykrobe predictor* on the *S. aureus* validation set (St_B1), treating the consensus phenotype as gold standard except for trimethoprim, which Phoenix does not test, where the disc test was used as truth. FN: False negative calls. R: total number of resistant samples. FP: false positives. S: total number of susceptible samples. VME: very major error rate (false negative rate), only shown where R>10. ME: major error rate (false positive rate), only shown where S>10. Error rates meeting the FDA requirements are in bold.

| Drug | FN(R) | FP(S) | VME (95% CI) | ME (95% CI) |
| --- | --- | --- | --- | --- |
| Penicillin | 0 (377) | 11 (94) | **0.0% (0.0%-1.0%)** | 11.7% (6.0%-20.0%) |
| Erythromycin | 1 (79) | 2 (392) | **1.3% (0.0%-6.9%)** | **0.5% (0.1%-1.8%)** |
| Ciprofloxacin | 3 (65) | 1 (406) | 4.6% (1.0%-12.9%) | **0.2% (0.0%-1.4%)** |
| Methicillin | 0 (54) | 0 (417) | **0.0% (0.0%-6.6%)** | **0.0% (0.0%-0.9%)** |
| Fusidic acid | 0 (41) | 4 (430) | **0.0% (0.0%-8.6%)** | **0.9% (0.3%-2.4%)** |
| Clindamycin | 0 (25) | 2 (97) | **0.0% (0.0%-13.7%)** | **2.1% (0.3%-7.3%)** |
| Tetracycline | 0 (17) | 2 (454) | **0.0% (0.0%-19.5%)** | **0.4% (0.1%-1.6%)** |
| Rifampicin | 0 (5) | 0 (466) | N/A | 0.0% (0.0%-0.8%) |
| Gentamicin | 1 (3) | 0 (468) | N/A | 0.0% (0.0%-0.8%) |
| Mupirocin | 0 (2) | 0 (348) | N/A | 0.0% (0.0%-1.1%) |
| Trimethoprim | 0 (1) | 1 (188) | N/A | 0.0% (0.0%-2.9%) |
| Vancomycin | 0 (0) | 0 (471) | N/A | 0.0% (0.0%-0.8%) |

The equivalent plot for false positive calls is shown in Fig. 2d. For all drugs except penicillin and methicillin, all methods have low false positive rates, below the FDA threshold of 3%. All methods had unacceptably high error rates for penicillin (11.7% for *Mykrobe*, 15.1% for disc, 16.0% for Phoenix). However, it is known that for penicillin, phenotyping methods may under-call resistance^17^-^19^, and so the apparent high false positive rate is likely to be artefactual – i.e. under-calling resistance by disc, Phoenix and nitrocefin might lead to an (incorrect) consensus susceptible call. Indeed in Gordon *et al* (2014)^3^ it was found that for some samples with weak beta-lactamase activity (exhibited by a very slowly developing nitrocefin test), resistance was not called by one of disc/Phoenix. *Mykrobe* had an acceptable false positive rate for methicillin of 0.0%, compared with disc (0.5%) and Phoenix (18.9%). This high false positive rate for Phoenix was unexpected; either both disc and Etest under-called resistance (they are both diffusion methods and might have correlated errors), or these were indeed false calls from Phoenix.

For comparison with a commercial software package, we also ran SeqSphere^6^ on the validation set, which made predictions for six drugs (methicillin, erythromycin, clindamycin, gentamicin, mupirocin and vancomycin) where resistance was gene-based. SeqSphere predicted all 471 samples to be resistant to erythromycin and clindamycin, so we excluded these drugs (after discussing with the author, D. Harmsen). Other results were broadly comparable to *Mykrobe predictor* and the phenotyping methods. See Supplementary Table 5 and Supplementary Figs. 4-5 for full results.

Finally, our strong prior expectation was that there would be limited within-sample diversity, due to blood culture followed by storage processes, and removal of contaminated samples (see Methods). *Mykrobe predictor* confirmed this expectation and made no minor calls. However we noted with interest that for the four samples where *Mykrobe* made false positive (major) resistant calls that were not made by Disc or Phoenix, re-running the disc test resulted in contradictory results. Two changed to resistant (ciprofloxacin, erythromycin) and 2 produced heteroresistant phenotypes (erythromycin, tetracycline, see Fig. 3). This behavior is consistent with a disc test presented with mixed strains or with variable plasmid loss (which would explain the 3 erythromycin/tetracycline results). All 4 samples had low levels of chromosomal diversity (between 12 and 25 “heterozygous” SNPs, Supplementary Methods), ruling out contamination unless by a closely related strain.

**Fig. 3.**
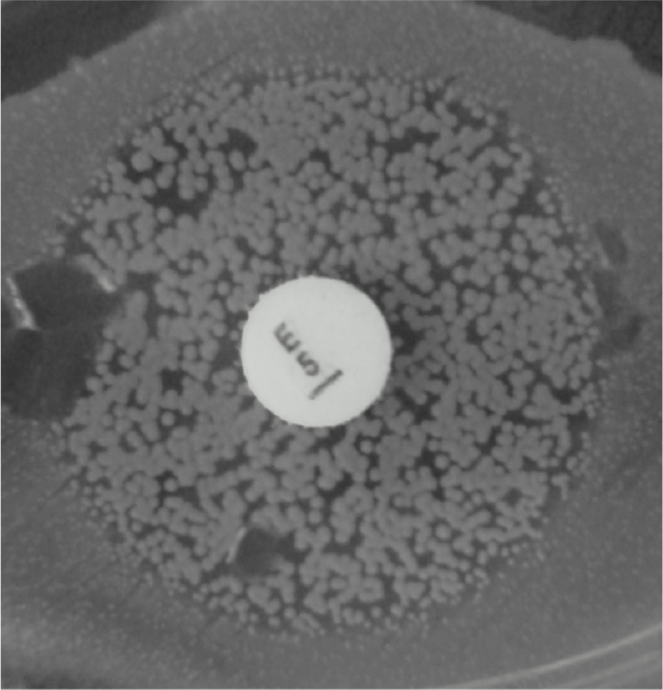
Photograph of heteroresistant phenotype seen on re-running Erythromycin disc test on a sample where *Mykrobe* had called a false positive (resistant) that neither Disc nor Phoenix had called.

### Simulating minor infections with empirical data

In order to determine the power of our method to detect minor resistant populations, we took 450 samples from the *S. aureus* dataset St_B1 that all had at least 100× mean sequencing depth of coverage across the genome, and subsampled them randomly to precisely 100×. We then took 1000 random pairs of samples from this set, and for each pair, combined subsets of their reads so as to create 27 different mixtures with ratios ranging from 1:99 to 99:1 – we call this Simulation We ran the *Mykrobe predictor* on all 27,000 mixed infections. The results are shown in Fig. 4a. Our method has greatest power to detect resistance to tetracycline and erythromycin, mediated by genes on multi-copy plasmids, reaching 90% by the time the population frequency is 2% for tetracycline, and 3% for erythromycin. For other drugs, detection power exceeds 90% once the subpopulation exceeds 10% frequency. This sensitivity did not come at the cost of loss of specificity - there were only 20 false positive calls out of 189,000 (27,000 mixtures × 7 drugs).

**Fig. 4.**
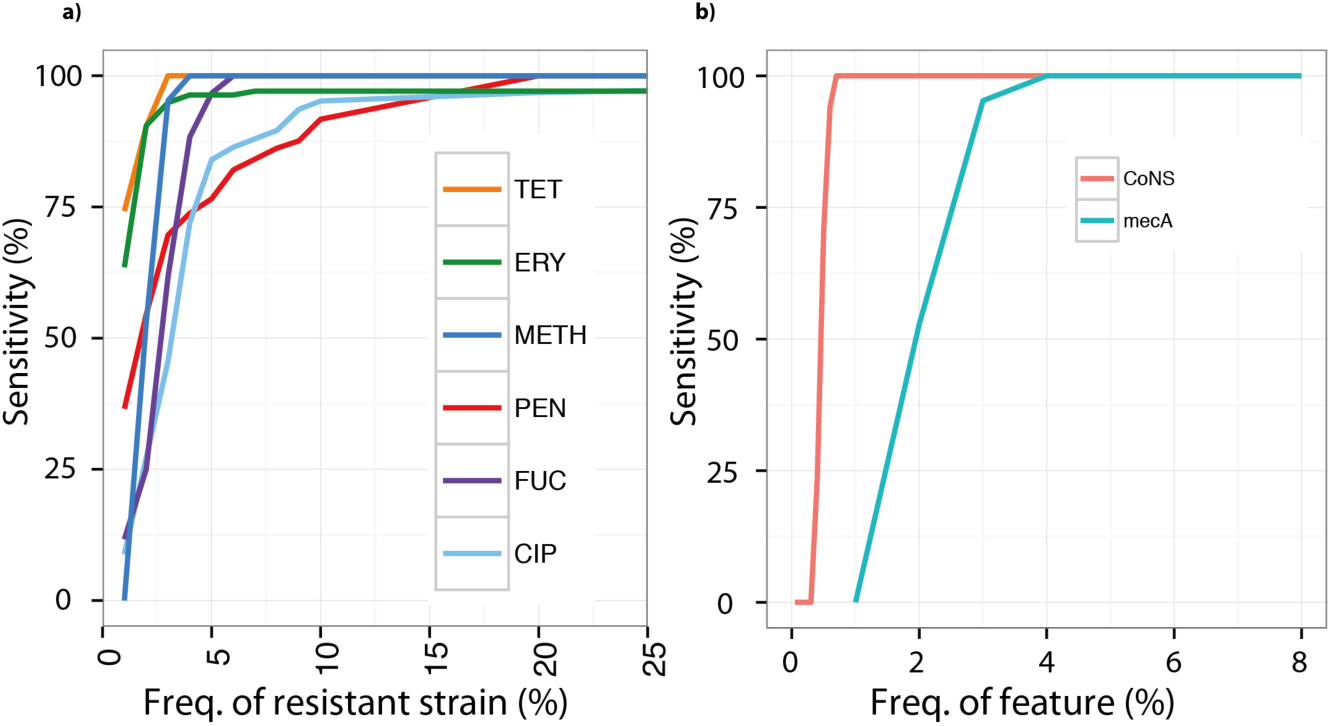
a) Simulation 2: power to detect minor resistant population in simulated mixtures of empirical data. As above, we do not estimate false negative rates for drugs where we have less than 10 resistant samples, as confidence intervals would be unreasonably large. Power is greatest for the drugs where resistance genes reside on multi-copy plasmids, erythromycin and tetracycline. Abbreviations: Tet: tetracycline, Ery: erythromycin, Meth: methicillin, Pen: penicillin, Fuc: fusidic acid, Cip: ciprofloxacin. b) Power to detect low frequency coagulase-negative species (red, Simulation 1, described above) is consistently higher than power to detect *mecA* (blue, Simulation 2), which causes methicillin resistance in *S. aureus*. Thus the risk of mis-calling MRSA due to detecting *mecA* from undetected contaminating coagulase-negative species is limited.

One goal for *Mykrobe predictor* was to minimize the mis-calling of MRSA caused by *mecA* from undetected contaminating CoNS. We therefore used the simulated mixtures of *S. aureus* and CoNS (Simulation 1), to estimate the discovery power for low frequency CoNS species, and compared with that for low frequency *mecA* in Simulation 2 - see Fig. 4b. We were able to confirm that in these mixtures such miscalls were indeed unlikely – at 1% frequency, the estimated power to detect the presence of a CoNS species was 100% (red curve), but power to detect *mecA* was 0 (blue curve). Above 4% frequency, power to detect each was 100%.

### Virulence elements

Antimicrobial resistance is not the only medically relevant phenotype that might be revealed by sequencing. *S. aureus* has a large number of virulence elements which might prove valuable to genotype. As an example, we considered Panton-Valentine Leukocidin (PVL), a cytotoxin that kills leukocytes and is associated with tissue necrosis^20^. We tested for presence of the PVL genes *lukPV-S* and *lukPV-K* on 67 samples from an outbreak, and validated by comparison with PCR tests for the presence of these genes (23 negative and 44 positive). The results were 100% concordant for all samples.

### Mycobacterial species identification

The second species we applied the *Mykrobe predictor* to was *M. tuberculosis*. We focused on the use-case where a sample undergoes Mycobacterial Growth Indicator Tube (MGIT) culture, after which the most likely species present is a Mycobacterium.

We combined datasets MTBC_A1 and Myco_SRA to generate probes (described in Methods) for detection of 4 MTBC species (*M. tuberculosis*, *M. bovis*, *M. africanum, M. caprae*) and 40 NTM species including *M. abscessus*, *M. avium* and *M. intracellulare*. If present, co-infection with both MTBC and NTM, which is known to occur^21,22^, would be reported. We also use SNPs defined in Stucki (2012)^23^ to identify lineage within the MTBC. In terms of desired error-profile, the main aim would be to minimize misclassifying a MTBC as a NTM, or vice-versa. Misidentifying species within MTBC has limited impact on choice of treatment, except that *M. bovis* is known to be intrinsically resistant to pyrazinamide and some Bacille Calmette–Guérin (BCG) strains are known to be resistant to isoniazid. We then evaluated species prediction on the union of datasets MTBC_A2 (N=1157) and Myco_Retro (N=147), where species had been identified by Hain assay, showing results in Fig. 5a. No samples were misclassified between MTBC and NTM, but there were 4 *M. africanum and 2 M. tuberculosis* samples that were only resolved to MTBC, and 1 *M. tuberculosis* sample misidentified as *M. africanum*. See Supplementary Table 6 for the full results table. Finally we tested our identification of the lineages as defined by Gagneux *et al* in^24^ by comparing with the lineage as identified by their own tool, KvarQ^11^ and found 100% concordance.

**Fig. 5.**
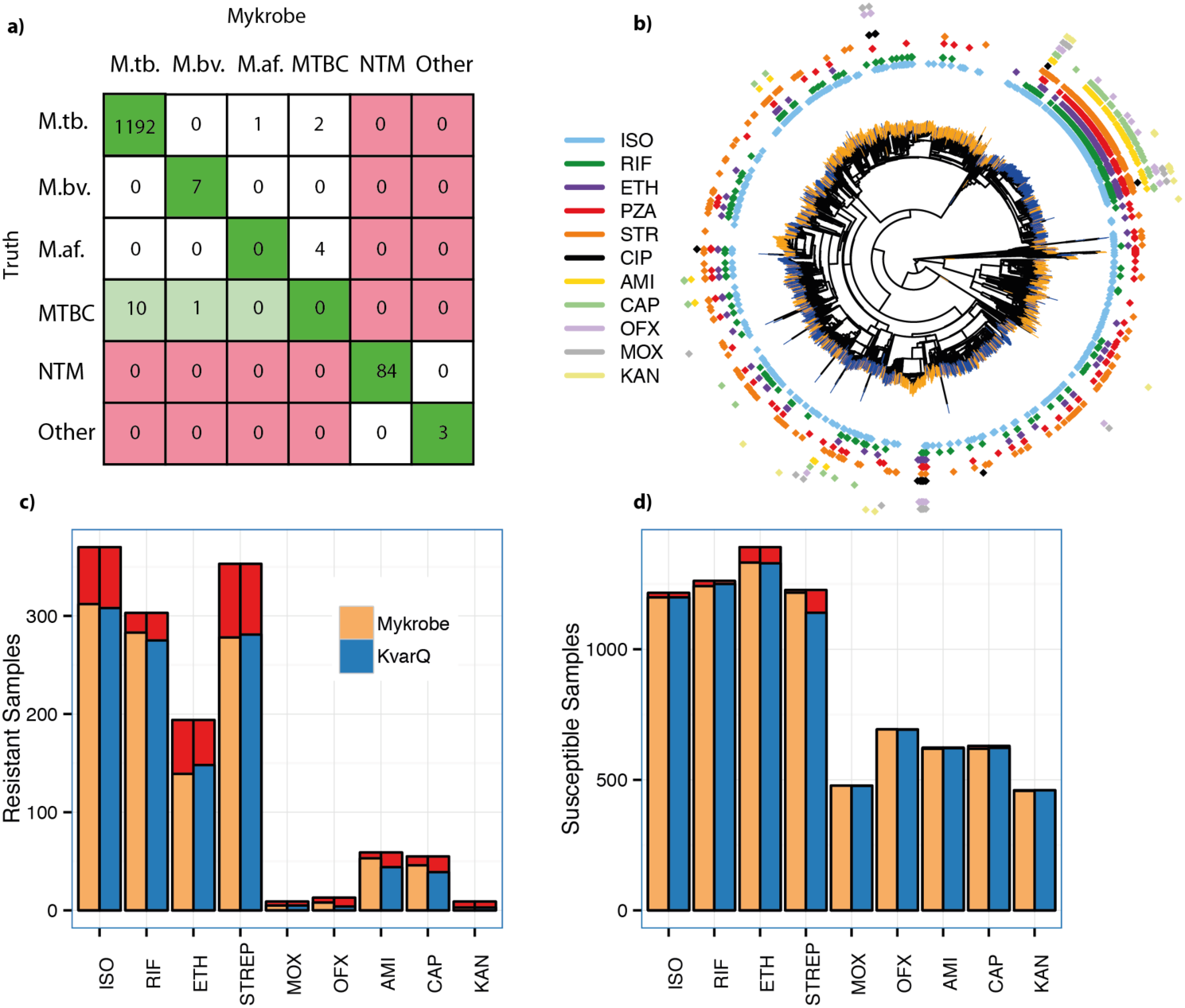
Species predictions for Mycobacteria and resistance predictions for *M. tuberculosis* complex. a) Species classification results on a validation set (MTBC_A2 + Myco_Retro). Colours indicate misclassifications between NTM/MTBC (red), concordance with “truth” (dark green), or greater resolution from *Mykrobe* than PCR (light green). Abbreviations: M.tb.: *M. tuberculosis*, M.af.: *M. africum*, M.bv.: *M. bovis*. b) Phylogeny of *M. tuberculosis* complex samples with phenotype data, with tips marked orange/blue to show if sample in training set (MTBC_A) or validation set (MTBC_B), and showing drug resistance in concentric rings around outside. Resistance exists across the phylogeny, especially against isoniazid (light blue), with a clustering of multi-drug resistance in the Beijing lineage. c) Proportion of resistant *M. tuberculosis* complex samples called as resistant for *Mykrobe* (yellow) and KvarQ (light blue) compared with phenotype - false negatives in red, d) As c) but for calling phenotypically susceptible samples as susceptible – false positives in red.

### M. tuberculosis – resistance predictions

We use a “training” dataset MTBC_A of 1920 samples from Oxfordshire, Birmingham, Sierra Leone and South Africa purely to fit the frequency parameter for the *Mykrobe predictor* minor-resistant model (see Methods), and a separate dataset (MTBC_B) of 1609 further samples from Uzbekistan, Germany, South Africa and the UK to validate. These samples were all collected for an independent study^25^ on the discovery of mutations predictive of resistance.

Figure 5b shows a phylogeny of these samples, with the sample membership of training/ validation set marked at the leaves of the tree in orange/blue. The validation set does show some clustering within the phylogeny, due to the large number of samples from Uzbekistan in the validation set.

Our understanding of the genetic basic for resistance in MTBC is not complete. Common resistance mutations are on commercial line probe assays, and explain approximately 85-95% of observed resistance to the two primary first line drugs (isoniazid, rifampicin)^26^-^28^. These assays have lower sensitivity for the fourth first-line drug (ethambutol) and second-line drugs^29^, and do not attempt to predict resistance for the third first-line drug (pyrazinamide), which is poorly understood. We built a panel of resistance mutations based on the Hain and AID line probe assays, with a small number of additional mutations from the literature (see Methods for details). For comparison with a method using a similar panel but without minor calls, we also ran the KvarQ tool^11^, which uses kmer recovery to detect alleles.

Figure 5, panels c,d show the proportion of resistant and susceptible samples that were called correctly for each drug. As expected, for first-line drugs rifampicin, isoniazid and ethambutol the two methods (*Mykrobe* and KvarQ respectively) have similar power to detect resistance (93.4%, 84.3%, 71.6% versus 90.8%, 83.2%, 76.3%) and similar false positive rates (1.6%, 1.4%, 4.2% versus 1.0%, 1.4%, 4.5%) - in line with expected performance of the Hain assay (Supplementary Fig. 6-7). We discuss below the higher number of false resistant calls in rifampicin made by *Mykrobe*.

Fewer samples were phenotyped for second-line drugs, but *Mykrobe* had noticeably higher sensitivity for amikacin and capreomycin (89.9% and 83.6% respectively) than KvarQ (74.6% and 70.9%), see below. There were very few false calls for second-line drugs for either method, except for a high (7%) error rate for KvarQ for streptomycin. See Supplementary Tables 7-10 for the full results on training and validation sets.

### Rifampicin false positives

On examination, we found the 20 false rifampicin-resistant calls may reflect limitations of culture-based phenotyping, rather than lack of power in *Mykrobe*. We show in Fig. 6a the estimated within-sample frequency of these calls. Firstly, one subset (n=8) had within-sample frequency of 5-10%, and recent tests suggest standard DST can fail to detect resistance at frequencies below 10%^30,31^ – thus these may actually be false negatives by the DST. The remainder (n=12) were all major calls, and were either mutations that have been shown to slow growth, or are at very nearby sites^32^-^34^. The specific mutations are Gln-429-His (n=1), Leu-430- Pro (n=6), Ser-450-Leu (n=3), Ser-450-Trp (n=1), Leu-452–Pro (n=1) in *rpoB*, the RNA polymerase gene. However, since the proportion method used in *M. tuberculosis* susceptibility tests fundamentally measures growth rate as a proxy for resistance^35^, this can lead to false susceptible DST calls for samples with these mutations^31,36^-^38^. In short, false positive calls from *Mykrobe* may actually be resistant *in vivo*, but called susceptible by gold-standard DST due to the nature of the test.

**Fig. 6.**
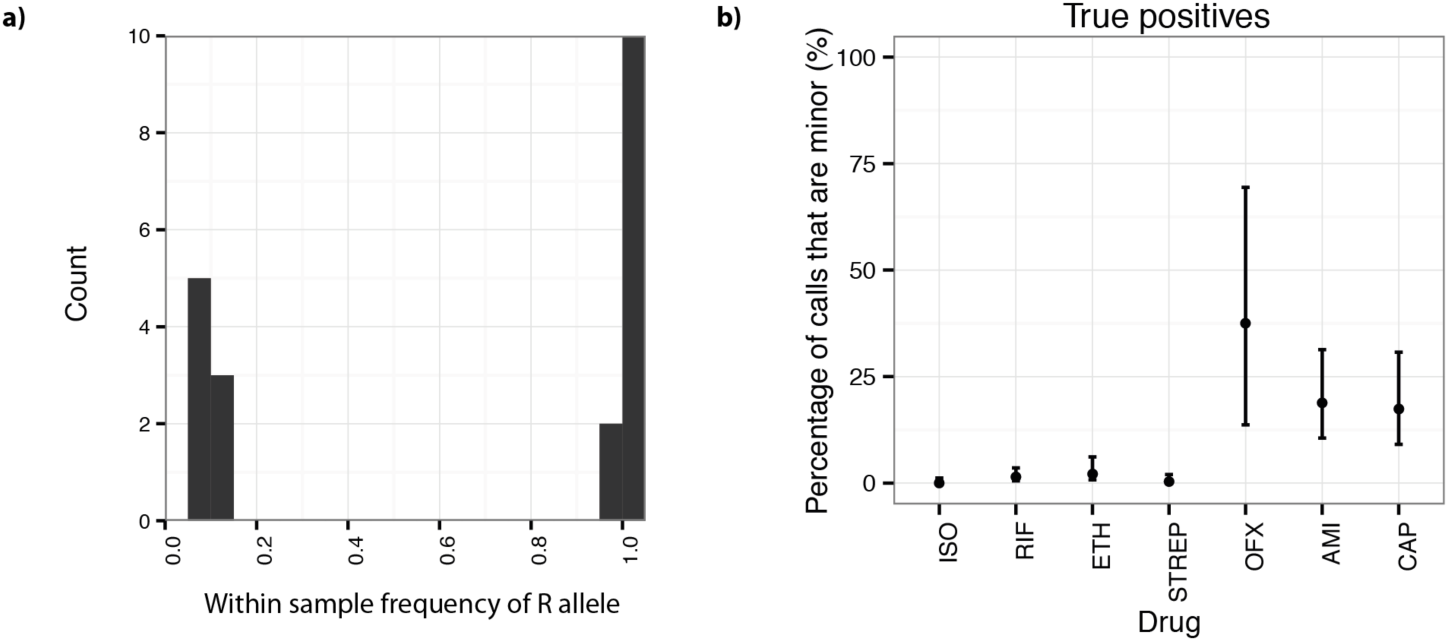
a) Estimated within-sample frequency of Rifampicin false positives, b) Percentage of true positive resistant calls due to the minor-population model; confidence intervals are calculated using the Clopper-Pearson interval. Drugs with less than 10 resistant samples excluded, to avoid overly large confidence intervals. For aminoglycosides and quinolones, detecting minor populations increases power to predict phenotypic resistance by greater than 15%. Abbreviations: ISO: isoniazid, RIF: rifampicin, ETH: ethambutol, STREP: streptomycin, OFX: ofloxacin, AMI: amikacin, CAP: capreomycin.

### Minor resistant population calls for M. tuberculosis

*Mykrobe predictor* made 75 minor-resistant calls across the 9 drugs and 1609 samples in the validation set. Whole genome analysis of these samples (see Methods) found a median of 16 heterozygous sites per sample, consistent with mixed infections (local transmission or in-host evolution)^39^, although we cannot exclude the possibility of contamination with a closely-related strain.

We show in Fig. 6b the proportion of true positives due to minor-resistant calls, showing a clear demarcation between first and second-line drugs. Power to predict phenotypic resistance to second-line drugs amikacin and capreomycin was significantly increased by the detection of minor populations, from 69% to 84% for capreomycin, 73% to 90% for amikacin and (although here the numbers were small (N=13)) 38.5% to 61.5% for ofloxacin. The increase in sensitivity from including minor-resistant predictions did not cause a loss of specificity, except for rifampicin, where 8 of the 20 false positives were due to minor calls, increasing the false positive rate from 1.0% to 1.6%. However, as discussed above, these calls may have clinical value despite discordance with phenotype. Indeed there is evidence that these mutations may be associated with poor outcome^40,41^.

### Nanopore sequencing

We tested *Mykrobe predictor* on data from the Oxford Nanopore Technologies MinION single-molecule sequencing machine. Since the per-base error rate is high (between 10-30% per base, depending on whether the molecule has been sequenced in one or two directions, termed “1d reads”and “2d reads” respectively), we modified the *Mykrobe predictor* to expect an error rate of 10%, and to detect a gene if it recovered 50% (rather than 90%). We took a multi-drug resistant *S. aureus* isolate from a clinical sample taken in 2014, and sequenced it with both the Illumina MiSeq and a MinION (see Methods for details) and ran the *Mykrobe predictor*. The MiSeq run took 24 hours and produced, after cutting reads at bases with quality below 10, 368x of 122bp reads. The MinION run took 24 hours and generated 39x of “2d” reads, with min/mean/max length 113bp/4.7kb/48kb. In both cases, *Mykrobe* correctly predicted the sample was resistant to penicillin, methicillin, gentamicin, trimethoprim, erythromycin, ciprofloxacin and clindamycin, and susceptible to fusidic acid, rifampicin, tetracycline, vancomycin and mupirocin. All of the resistance calls were due to detection of genes, except for ciprofloxacin where a Ser->Leu mutation at position 84 in the gene *gyr* was detected. No false positive resistance SNPs were called. Furthermore, by truncating the MinION output file we showed these results could be obtained with just 8 hours of sequencing. We conclude that it should be feasible to apply *Mykrobe* to this new strand-sequencing technology.

### Software performance and usability

*Mykrobe predictor* memory use is comparable with that typically used by a web browser such as Chrome with multiple tabs open, and CPU requirements are low; *Mykrobe* has been run on a Google Nexus 10 tablet, a Samsung Core Duos phone and a Raspberry Pi Model B. It scales across multiple CPUs on a multi-core server or on a compute cluster. We give in Table 2 some basic performance statistics for *S. aureus* (abbreviated Sa) and *M. tuberculosis* (abbreviated Mtb) data.

**Table 2.** Performance and feature comparison. Performance and feature comparison. We show elapsed time for one sample on a laptop (Macbook Air with 8GB RAM) and a server (Dell PowerEdge R820 with 32 cores, 1Tb RAM), and then for the entire *S. aureus* and *M. tuberculosis* validation sets. We ran KvarQ on 1 thread for ease of parallelization and comparison, as recommended by the authors. However we ran SeqSphere on 4 threads because to use 1 would have taken a prohibitively long.

|  | Mykrobe (Sa) | SeqSphere (Sa) | Mykrobe (Mtb) | KvarQ (Mtb) |
| --- | --- | --- | --- | --- |
| RAM | 100Mb | 8Gb | 100Mb | 30Mb |
| Time/sample on laptop | 1.5 mins | - | 2.75 mins | 40 mins |
| Time/sample on server | 44 sec | 19 mins | 47s | 23 mins |
| CPU time (Mtb validation set) | - | - | 1 day | 30 days |
| CPU time (Sa validation set) | 5.8 hours | 6.2 days | - | - |
| Speciation of clinical samples | Yes | No | Yes | No (MTBC only) |

## Discussion

Rapid determination of antimicrobial resistance profile is of critical importance to patient care for many serious bacterial infections and has wider implications for determination of treatment protocols and national surveillance. We have developed a generic framework, extensible to many species, called the *Mykrobe predictor*, which can identify species, resistance profile and other genomic features such as virulence elements and phylogenetic lineage, within 3 minutes on a standard laptop. We have provided two implementations, for *S. aureus* and *M. tuberculosis*, validated extensively against clinical gold-standards. Our results for *S. aureus* (overall sensitivity and specificity above 99%) are comparable with or better than phenotyping methods (BSAC disc test, Phoenix). For *M. tuberculosis*, specificity is high (98.5%) and sensitivity of 82.5% matches the line probe assays from which our resistance panel was constructed, but still is below that of the gold-standard of DST based on Lowenstein-Jensen culture. However, as new resistance-causing mutations are determined, *Mykrobe predictor* can be updated and tested extremely easily, and unlike a line-probe assay or the automated Xpert-Mtb/Rif (Cepheid) assay, is unaffected by size of resistance panel.

In terms of time from bedside to result, for *S. aureus*, using latest Illumina MiSeq reagents with 16.5 hour sequencing run, a sequencing-plus-*Mykrobe* workflow would be approximately 4.5 hours faster than the current clinical workflow (see Supplementary Fig. 8), taking 31.5 hours (although in this large retrospective study we used a HiSeq for higher throughput). For *M. tuberculosis*, most clinical isolates become MGIT positive within two weeks; if sample preparation and sequencing is completed within 2 days of positivity^42^, one can get results in 2 weeks, rather than approximately 6 weeks for standard DST.

For both species, the major time bottleneck is culture. Improvements in sample preparation and sequencing from populations with low quantities of pathogen DNA and/or high quantities of other bacterial DNA, will result in shorter culture times and less biased sampling of the within-host population. Such improvements are not possible for traditional susceptibility tests, as they rely fundamentally on the input sample being clonal and of a fixed inoculum size. We have demonstrated that simple minor resistant infection detection, assuming at most 2 strains are present, can be achieved in a robust and automated fashion. We are unaware of any other resistance prediction tool that allows this. In our study, we specified for *M. tuberculosis* a 20% frequency for the minor resistant model, and found that 4.7% (75/1609) of our validation samples had minor resistant populations. However, we chose 20% purely in order to give resistance predictions that matched culture-based DST; dropping the value to 10% would have called a further 19 samples (median coverage 127x) with minor resistant populations (median frequency 7.5%), but which corresponded exclusively to DST-susceptibility. Even after culture, these low frequency resistant alleles are clearly present in an appreciable fraction (94/1609) of clinical samples but do not survive culture-based DST. Large-scale collection of this information that *Mykrobe predictor* provides, combined with treatment and patient outcome information, may enable us to determine whether they are of clinical consequence despite failing DST, as has been suggested for some rifampicin resistance mutations^40,41^.

For TB, initial treatment is with the preferred set of 4 “first-line” drugs until it is identified that a drug-resistant strain is present, when treatment is changed to the less-effective, more complicated to administer (injectable rather than oral) and more toxic “second-line” drugs. The spread of MDR (multi-drug resistant) TB^43^, defined as resistant to the key first-line drugs rifampicin and isoniazid, places greater pressure to use second-line drugs, and therefore also to be able to detect resistance to them. Heteroresistance to second-line drugs has been previously reported^44^-^46^ as a matter of concern, consistent with our finding that minor population detection gives *Mykrobe* increased power for predicting resistance to amikacin, capreomycin and ofloxacin.

Recognizing that lack of bioinformatics expertise is a barrier to clinical adoption, we provide drag-and-drop Windows and Mac applications (see screenshot Supplementary Fig. 9-11) and a linux version that could for example enable a cloud service. Our demonstration that *Mykrobe* can work on low-specification hardware such as a mobile phone or Raspberry Pi is intended to enable future applications of resistance determination in the field, in low-resource settings with no internet access. Along these lines, the advent of portable single molecule sequencing machines which deliver long-read information in real-time will change the face of clinical microbiology. The ability to sequence a single sample removes the need to batch samples until an Illumina MiSeq or HiSeq sequencing run is justified, reducing bedside-to-treatment time, and the long reads could provide vital information on mixed infection composition. Our N=1 test of the Oxford Nanopore MinION machine offers little more than proof-of-principle, but we were struck that with only 13x coverage we were able to get fully concordant results for both gene and SNP-driven resistance, without the high error rate causing any false SNP calls. Since *Mykrobe* can test the de Bruijn graph and assess confidence of resistance/susceptibility approximately an order of magnitude faster than MinION reads arrive, this could be done as the reads come in from the machine, enabling a real-time decision to be made as to whether to stop sequencing. In this case we could have stopped after 8 hours of sequencing, half the time needed for an Illumina MiSeq run.

In terms of the path to clinical use, the next steps for the *Mykrobe predictor* for *S. aureus* and *M. tuberculosis* are further clinical testing, and then obtaining regulatory approval. We will be running *Mykrobe* in parallel with the clinical workflow in hospitals in Oxford, Leeds and Brighton for 3 months this year, as part of a Health Innovation Challenge Fund project. It is relatively simple to extend the *Mykrobe predictor* to other species by using a panel of known sites and genes as we did in this study. However more generally, for a species where resistance mechanisms are poorly characterized, one would need to use a large training set with both whole-genome sequence data and phenotype information for hypothesis-free discovery of causal mutations or purely predictive markers. Some such studies have been done^47,48^, and we expect many more.

There are two main limitations to the current implementation. First, we suspect that incorporating a more general model of mixtures, rather than simply major/minor clones, will be of value when analyzing *M. tuberculosis* samples direct from sputum. Secondly, for *M. tuberculosis* our sensitivity (82.5% across all drugs) is low compared with traditional DST, and completely excludes the first-line drug pyrazinamide since known mutations are poorly predictive. This issue, shared by all molecular assays, can only be resolved by large-scale sequencing and phenotyping studies.

## Methods

### Study Design

The objectives of this study were:

a. To show that our software program could deliver automated antimicrobial resistance predictions for 2 species given a *pre-specified* genotype-to-drug-resistance mapping. The limitations of the pre-specified mapping would place an upper bound on sensitivity – for *S. aureus* that upper bound was above 99%, but for *M. tuberculosis* we followed the HAIN and AID assays, expecting sensitivity of around 82%. To achieve this, we used independent training and validation sets previously obtained in other studies^3,25^. For both species, the number of samples with resistant phenotypes was limiting, and we only estimated false negative rates where there were sufficiently many (>10) resistant samples in the validation set, reporting confidence intervals calculated using the Clopper-Pearson interval.
b. To handle contamination and mixture issues seen in clinical samples. We used independent datasets for design of probes and validation. In order to include some realistic sampling of species, the validation set for mycobacteria included the set Myc_Retro consisting of all samples sent to the laboratories at the Oxford John Radcliffe Hospital between 2 June 2013 and 29 January 2014.
c. To observe whether minor resistant population detection could increase predictive power of phenotypic resistance without compromising specificity in the datasets collected.

All datasets used are described in Supplementary Fig. 2, and Supplementary Table 1 with full sample information in Supplementary Data Files 1-2.

### Phenotyping and sequencing of S. aureus datasets

Initial phenotyping of the training and validation sets was described in detail in Gordon et al (2014)^3^. The training set was phenotyped using either the Vitek automated system (bioMerieux) or the Stokes method disc diffusion^49^, whereas all validation samples were phenotyped using two methods: a BSAC disc test^14^ the Phoenix Automated Microbiology System (BD Biosciences, Sparks, MD, USA). For trimethoprim only disc testing was performed.

We removed 6 samples from the training dataset and 20 samples from the validation set that were contaminated (See Supplementary Methods for details). All samples where there was discordance between Phoenix and Disc for any drug in Gordon et al^3^ were rerun on Phoenix (for all drugs) and previous results from Gordon et al^3^ were discarded.

Samples were sequenced on Illumina HiSeq 2000 platform, with mean read length (after cutting reads at bases with quality score below 10) of 87 bp and mean depth of 87, as described in^3^. We double-checked, and by chance the previous two numbers are indeed both, independently, 87.

### Species identification in general

We developed a system for designing a hierarchy of markers (contigs) which first separated two phylogroups (*S. aureus* from coagulase-negative Staphylococci, or MTBC from NTM), and then identified species present within a phylogroup. We first build a Bruijn graph by pooling several hundred species from both phylogroups, and pull out all unique and unambiguous contigs (“unitigs”) and calculate the frequency of each contig in each phylogroup. We choose the most highly differentiated contigs to form marker panels to distinguish the groups. This process could then be run again to find contigs informative at the species level.

### Species id for S. aureus

The above process was applied to 731 Staphylococci in training set St_A, using Cortex^13^ with kmer-size 15, to produce probes (contigs) for phylogroups (*S. aureus* versus coagulase negative staphylococci) and species (*S. aureus*, *S. epidermidis*, *S. haemolyticus*, other coagulase-negative species. We assume a positive Gram stain has been obtained, and use 33 alleles of catalase gene (Supplementary Table 11) to confirm presence of staphylococci. Proportion of probe panel in each training sample (“recovery”) was plotted, and extreme outliers were ignored as possible errors in the SRA metadata; detection thresholds were chosen based on recovery in the training set: 90% for *S. aureus,* 30% for *S. epidermidis* and *S. haemolyticus,* 10% for other Staphylococcus and 20% for the catalase gene.

### S. aureus resistance panel

Starting from the variant catalogue introduced in Gordon *et al*.^3^ we made the following alterations. B434N in *fusA* was changed to D434N. Q456K in *rpoB* was removed as Q was not the amino acid at position 456 of the referenced *rpoB* gene. We added *rpoB* N474K from Villar (2009)^50^. We also considered the 10 novel mutations reported in Dordel (2014)^51^. We found 3 of the mutations (PBP1 H499Y, PBP2 T31M, PBP2 D156Y) in our derivation set. These were found in samples phenotypically susceptible to methicillin so none of the variants from this paper were included in the final catalogue. Variants that changed predicted MIC but did not confer resistance on their own were not included. For resistance genes we took all versions/alleles of the gene from NCBI that were not explicitly annotated as existing in a susceptible strain, and did not have stop codons. The full list of chromosomal mutations, genes, accession id’s can be found in Supplementary Tables 12-14.

### Data structures for genotyping

We implemented two versions of *Mykrobe predictor*. The first builds a whole genome de Bruijn graph of the sample, and then takes the intersection of this with the de Bruijn graph of all (alleles of) genes and mutations on different genetic backgrounds (the “target graph”). This requires approximately 300 Mb of RAM for a typical Illumina dataset, but could in principle grow for very large datasets. In order to control memory use, the second approach builds the target graph first, and then only loads sample data that intersects it, reducing RAM use to 100Mb. Both methods give absolutely identical results, and we ran all analyses for this paper using the second approach.

### Resistance calling at mutations

We use three competing models: pure susceptible, minor resistant (frequency=10% for *S. aureus*, 20% for *M. tuberculosis*), major resistant (we used frequency=75%, but we expect values from 60%-100% would all result in identical model choice). In this and subsequent sections, a subscript MAJ, MIN or S refers to the Major Resistant, Minor Resistant or Susceptible models. We use the following uninformative priors:

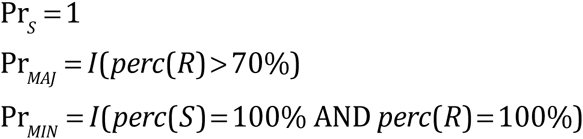

where *perc*(*R*) and *perc*(*S*) are respectively the percentage of the kmers in the resistant/susceptible alleles that are seen in the sample, and *I* is an indicator function. We use the following a simple Poisson model for the likelihoods for all 3 models

Susceptible model: Likelihood specified by Poisson coverage on S allele, plus errors driving both coverage loss on S allele and coverage on R allele.

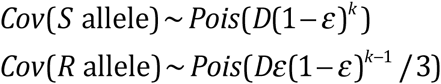

Major and Minor Resistant models: Poisson coverage on both alleles scaled by frequency:

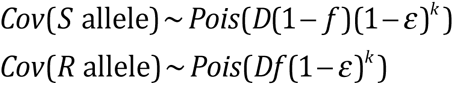

where *Cov*() is a function returning median coverage on an allele, *D* is the depth of coverage, *ε* is the per-base error rate, *k* is the kmer-size, and the frequency *f* of the resistance allele is 0.75 for the resistant model, and 0.1/0.2 for the Mixed model for *S. aureus*/*M. tuberculosis*.

The vast majority of mutations in the panel are amino acid changes in a gene, but some are promoter changes. When we refer to genetic background, we simply mean mutations present in the population within one kmer-length of the site of interest. For all mutations in the panel, we run through all genetic backgrounds, and if appropriate, all possible nucleotide changes that would generate the specified amino acid change, and find the highest coverage resistant allele and susceptible allele. These are then passed into the three models. The Maximum A Posteriori model is chosen.

### Resistance calling at genes

The expected proportion of kmers in a gene which are observed is

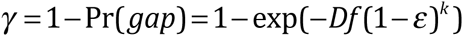

We use the following priors

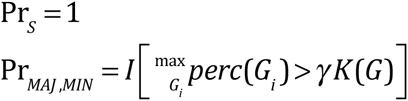

where each gene *G* has multiple exemplars *Gi* representing diversity of that gene, *I* is an indicator function, and *perc*() is a function returning the percentage of kmers present in the sample. *K* is a constant depending on the gene, set based on the levels of diversity seen in the training set:

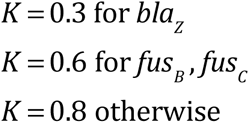

The likelihoods for Major and Minor models are the product of two factors: the probability of having the observed median coverage across the gene, and the probability of having observed *g* gaps. If *dpois* refers to the probability density of a Poisson distribution, and the observed median coverage across the gene is *m*, then the likelihood for the Major and Minor Resistant models are given by

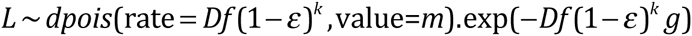

The likelihood for the Susceptible model is as in the mutation model – all coverage is due to errors.

### Species identification – M. tuberculosis

We chose to identify 4 MTBC species (*M. tuberculosis*, *M. africanum*, *M. bovis*, *M. caprae*) and 40 NTM species (including *M. avium*, *M. abscessus*, *M. intracellulare* – see Supplementary Methods for a full list). Marker panels were generated for MTBC and NTM as well as for individual species within those groups, using the same method as for staphylococci based on training set MTBC_A1 + Myco_SRA. The required detection threshold for each of the complex was set as 70% for MTBC, 25% for the NTM panel, and 30% for all species panels.

We also used the lineage-informative SNPs defined in Stucki (2012)^23^ to assign *M. tuberculosis* lineage: Beijing/East Asia, East Africa / Indian ocean, Delhi/Central Asia, European/American, West Africa 1 & 2, Ethiopian.

### M. tuberculosis phylogeny

We used the underlying phylogeny of samples in sets MTBC_A and MTBC_B, which was constructed using RAxML (version 8.0.5) using a GTRCAT model. For this study, we combined this tree with dataset membership and phenotypic resistance metadata using the R APE package^52^ to produce Fig. 2b.

### Resistance panel – M. tuberculosis

We used a panel of MTBC resistance variants from the HAIN^28^, Cepheid^53^ and AID^54^ assays supplemented by others from the literature ^55^-^57^ see Supplementary Table 15. All possible SNPs that would account for amino acid or DNA variant associated with resistance were introduced on multiple susceptible backgrounds. These backgrounds were selected as follows. Two samples were chosen from each of the 6 *M. tuberculosis* lineages. For those samples, paired-end reads were mapped by Stampy (version 1.0.17)^58^ to the H37Rv (GenBank NC000962.2) reference genome. SNP calls were made with SAMtools^59^ mpileup (version 0.1.18), requiring a minimum read-depth of 5 and at least one read on each strand. We looked for variants in the 20 bases on either side of each resistance mutation in each of those 12 samples – these, along with the reference, defined a set of genetic backgrounds. Since the underlying panel is almost identical, we expect *Mykrobe* to perform equivalently to the HAIN test. Comparing on MTBC_A1+MTBC_A2, *Mykrobe* and HAIN have similar power to detect resistance (85.5%, 94.1%, 76.5% versus 85.5%, 93.5%, 76.5%) for first-line drugs rifampicin, isoniazid and ethambutol respectively - see Supplementary Figs. 5-6.

We chose an underlying frequency of 20% for the minor resistant model as this gave appropriately low false positive rates when comparing with phenotypes in the training set (table S7).

### Software

The *Mykrobe predictor* software is available (open-source) at www.github.com/iqbal-lab/Mykrobe-predictor for academic and research use only under a license from Isis Innovation, the technology transfer company of the University of Oxford, that is free for non-commercial use. We provide a Linux command-line version and desktop “drag-and-drop” applications for 64-bit Windows and Mac OS X (see screenshot Supplementary Fig. 9-11).

## Acknowledgments

We would like to thank Michel Doumith and Angela Kearns for helpful discussions, Sebastian Gagneux and Andreas Steiner for help running KvarQ, and Dag Harmsen for help running SeqSphere. We are also grateful to the MinION Access Program from Oxford Nanopore Technologies which enabled us to trial their new sequencer. We acknowledge funding from UK Clinical Research Collaboration (Wellcome Trust [grant 087646/Z/08/Z], Medical Research Council, National Institute for Health Research [NIHR grant G0800778]), NIHR Oxford Biomedical Research Centre, NIHR Oxford Health Protection Research Unit on Healthcare Associated Infection and Anti-microbial Resistance, EU FP7 Patho-Ngen-Trace (FP7 - 278864- 2). ZI and DJW were funded by two Wellcome Trust/Royal Society Sir Henry Dale Fellowships (grants 102541/Z/13/Z and 101237/Z/13/Z respectively). PB was funded by a Wellcome Trust PhD studentship, and SE was funded by an MRC funded prize studentship to the Nuffield Department of Medicine, University of Oxford. DWC and TEAP are NIHR Senior Investigators. GM was funded by grant 100956/Z/13/Z from the Wellcome Trust.

## Author contributions

Designed the study: ZI, GM, PB. Wrote the paper: ZI. Designed and wrote core software: PB, ZI. Windows/Android support: BH, PB. Performed computations/data analyses: PB. Determined clinical requirements: NCG, TMW, TEAP, DWC, LJP, DHW. User interface: SH. Reviewed drafts of paper: GM, ASW, DHW, DWC, LJP, NCG, TMW. Phylogeny construction: TG, SE, PB, DJW. Phenotyping: LD, NCG. Sample preparation for MinION and MiSeq: LA, AAV. MinION runs: PP, MdC. Resources for MinION: DB, ZI, DWC. Lab resources for phenotyping and samples: RD, SN, TAK, SOV, NI, EGS. All authors reviewed final draft.

## Competing interests

None.

## Data and materials availability

All sequencing data is either already available in the European Nucleotide Archive, or in the process of submission, and will be available by publication. Accession identifiers (where known) along with phenotyping data are in the supplementary data files 1 and 2 (see below).

